# Class background reverses the effect of a polygenic index of cognitive performance on economic ideology

**DOI:** 10.1101/2022.12.12.520104

**Authors:** Rafael Ahlskog

## Abstract

Variation in political preferences is increasingly understood to stem from both environmental influences and genetics. Research has argued that a possible pathway for genetic effects on ideology is via cognitive performance, showing a genetic overlap between the traits. Yet, an unambiguous link between cognitive performance and economic policy preferences remains elusive, with results ranging from negative to positive effects on economic conservatism. In this study, I argue that this heterogeneity reflects an underlying gene-by-environment interaction. I depart from the assumption that cognitive performance, rather than being directly connected to a set of policy preferences, influences the capacity to correctly optimize those preferences. Combined with insights from standard models in political economics, this leads to the proposal that genetics associated with cognitive performance should cause more left-wing economic preferences if you grow up in relative poverty, but more right-wing preferences if you grow up in relative affluence. Utilizing variation in a polygenic index of cognitive performance within dizygotic twin pairs, coupled with unique register data on economic conditions for both the twins, their parents, and their childhood neighborhood, I show that the effect of the PGI on a finegrained measure of economic conservatism is zero on average, but indeed sizable and sign-discordant by class background. To my knowledge, this represents the first sign-discordant GxE finding for a socially relevant outcome, which has implications for future social science research using PGIs.

## Introduction

As was famously argued by Aristotle, man is by nature a political animal – we organize groups and societies and argue over the common good and the distribution of resources among various sectors in society. While it may be true that we share a common political core, we still differ tremendously on what we think the political arena ought to do and accomplish, with some favoring large amounts of taxation and redistribution while others would prefer as little state involvement as possible. Consequently, understanding differences in economic policy preferences and ideology lies at the heart of political science and political economics writ large, with explanatory models dating back as far as the inception of the social sciences (e.g. Marx 1887).

Traditional accounts have typically centered on the influence of immediate environmental factors and socialization.^1^ For example, the foundational model in political economics argues that anyone below median income should prefer more taxes and redistribution, while anyone above median income should prefer less (Meltzer and Richard 1981, Romer 1975). Others have emphasized intergenerational transmission in the form of parental socialization and class voting (Anderson and Davidson 1943, Evans 2000, van der Waal et al. 2007). In the last decade and a half, however, a growing literature has begun investigating the role of genetic factors in explaining differences in political behavior. A large number of studies, using a range of methods and designs, have now documented the non-negligible influence of genetics (e.g. Dawes et al. 2021, Hatemi et al. 2014, Oskarsson et al. 2022).

This has raised the question of what the character of these genetic effects really are. While our evolutionary environment was probably marked by the group dynamics of resource sharing and group competition (Bowles and Gintis 2011), modern mass political systems characterized by delegation of power, taxation and public spending did not exist. Hence, our behaviors on this arena are unlikely to have been directly shaped by evolutionary forces. Genetic effects on economic ideology must therefore be mediated by more proximal traits – so-called endophenotypes. Understanding genetic effects on ideology therefore requires that we find out what these endophenotypes could be.

The research problem addressed in this study stems from literature showing a) that there appears to be some genetic overlap between ideology and cognitive performance, indicating that some of the genetic effects on political preferences may be mediated by traits related to cognitive functioning (Oskarsson et al. 2014), and b) that the relationship between actual trait cognitive performance and economic ideology, paradoxically, is wildly inconsistent across studies (Jedinger and Burger 2022). I propose that to resolve this apparent paradox we need to approach this problem from a gene-by-environment interaction (GxE) perspective; that is, when the effect of a given genetic factor differs depending on the environment in which it is expressed.

This study, thus, is an attempt to both contribute to the understanding of the genetic etiology of economic ideology, and to clarify its relationship with cognitive performance. To accomplish this, I depart from traditional rational choice models in political economics as well as long-standing insights into the importance of early-life environment and the stickiness of policy preferences. On this basis, I argue that cognitive performance, rather than influencing policy preferences directly, should affect one’s capacity to correctly optimize those preferences – or in other words, to understand one’s political self-interest. This leads to the GxE hypothesis that genetics associated with cognitive performance should cause more left-wing economic policy preferences in poor families, but more right-wing ditto in rich families.

I test this proposed GxE effect by relying on within-family differences in a polygenic index (PGI) of cognitive performance, in a large sample of genotyped Swedish dizygotic (DZ) twins. This design makes sure that estimates for genetic effects have a causal interpretation. Furthermore, the outcome is a fine-grained measure of economic ideology based on a large survey battery of political preference items. I combine this with unique register-based data on socioeconomic factors for both the twins themselves, their parents, and their rearing environment, and a number of strategies to safeguard against confounding of the moderating effect.

The results show that class background not only moderates the effect size of the cognitive performance PGI, but indeed reverses it: among individuals from families in relative poverty, the PGI causes more left-wing economic views, but among their affluent counterparts it instead causes more right-wing economic views. On average, the PGI effect is in fact zero.

This study thus reinforces the view that the effects of cognitive performance itself are divergent and dependent on third variables. Instead of a simple direct effect on economic ideology, it most likely affects the antecedents of these preferences: one such antecedent is, arguably, the extent to which one is able to comprehend one’s class interest. In effect, genetics associated with cognitive performance can indeed help connect the dots between genetics and political outcomes, but, at least when talking about economic ideology, these effects can only be understood as environmentally contingent.

To my knowledge, this study represents the first robust finding of a so-called “lens”-type GxE effect (Domingue et al. 2020) for a socially relevant outcome, where not only the effect *size*, but even the effect *sign* differs, and where the average effect is zero. While the results in themselves should be of relevance for a wider audience interested in preference formation more generally, they therefore also underscore an important point about the methodology of using PGI’s as predictors of various social and behavioral phenotypes other than their target traits: given the likely complex interdependence between genes and environment, average observed genetic effects (including null results) may sometimes hide a substantial amount of theoretically highly interesting heterogeneity.

## Theory

Since the publication of a seminal twin study on the heritability of political preferences by Alford et al. (2005), a plethora of studies have documented a moderate to large amount of heritability for a variety of politically relevant phenotypes, such as self-reported ideological placement and economic and social attitudes (Hatemi et al. 2014), political participation (Dawes et al. 2014) and social trust (Oskarsson et al. 2012). In trying to find the mechanisms, or endophenotypes, for these proposed genetic effects, Oskarsson et al. (2014) finds that a range of ideological dimensions have substantial genetic overlap with cognitive ability. For example, the genetic correlation is shown to be 37% for redistributive preferences, 35% for immigration policy attitudes and 32% for foreign policy preferences. Genetic overlap, especially when the proposed endophenotype (i.e. cognitive ability) is more proximal to the underlying biology, is an indication that some of the genetic effect on the ideological outcome is mediated by that endophenotype.

This raises the question of what, in turn, the connection between trait cognitive performance and different ideological constructs is. As argued by among others Feldman and Johnson (2014), the minimum number of ideological dimensions needed to understand the structure of public opinion is two: economic ideology, which captures variation in attitudes related to taxes, redistribution and government intervention into the economy, and social ideology, which captures moral and cultural issues like abortion, gay rights, immigration etc. When it comes to social ideology, the connection to cognitive performance is rather clear in the literature, with the vast majority of studies showing a negative correlation between measures of IQ and social conservatism (e.g. Carl 2015, Deary et al. 2008, Eidelman et al. 2012, Ludeke and Rasmussen 2018, Schoon et al. 2010).

The connection between cognitive performance and economic ideology, however, is less than clear, with both positive and negative effect estimates in the literature. On the positive side, one study finds that intelligence predicts higher economic conservatism in the American National Election Study (Carl, 2015). Another study by Ludeke and Rasmussen (2018) finds that intelligence is uncorrelated with vote choice in both the US and Denmark, but that this lack of correlation merely obscures differential effects on social and economic ideology. Countering evidence is provided by studies finding that people with more free market-oriented preferences score worse on measures of cognitive biases (Sterling et al. 2016), while others find negative but insignificant correlations between measures of cognitive performance and economic conservatism (Saribay and Yilmaz 2017, Choma et al. 2019). More recently, a meta-analysis of the relationship finds no less than 19 studies that pass inclusion criteria, with point estimates of the correlation ranging from −.13 to .25 (Burger and Jedinger 2022). Furthermore – and most importantly – this meta-analysis concludes that the heterogeneity is so large that it must be explained by third factors rather than sampling error alone.

The premise for the following line of argumentation has thus been laid out: cognitive performance is a plausible candidate for an endophenotype connecting genes to economic ideology, and yet, some third factor is required to make sense of the disparate body of evidence on the effects of trait cognitive performance. In the following, I will argue that such a third factor is socioeconomic status, and more specifically class *background*. To connect the dots between class background, economic ideology and the genetics of cognitive performance, and in the end derive the proposed GxE hypotheses, we will start with a brief overview of the fundamental rational choice logic of policy preferences.

The foundational model of rational economic policy preferences is often condensed into a simple model with a flat tax rate and a lump sum transfer, whereby the state taxes all citizens a fixed *proportion* of their income, and gives back a fixed *nominal* sum to everyone (Meltzer and Richard 1981, Romer 1975). The predictions of such a model are that anyone earning less than the mean income should prefer complete redistribution, whereas anyone earning above the mean should prefer no redistribution whatsoever.

While this is a crude oversimplification of real-world political systems, the fundamental dynamics are robust: people with low incomes tend to be net-beneficiaries of redistribution, whereas the opposite is true for those with high incomes. In general, therefore, we should expect a clear socioeconomic gradient in degrees of economic conservatism. Such a gradient is also well supported in, among other things, the literature the class-basis of voting (Anderson and Davidson 1943). While many have argued that the impact of class based voting has declined over time (Clark and Lipset 1991, Manza et al. 1995, Evans and Tilley 2012), others have claimed that this supposed decline has been greatly exaggerated (Evans 2000, van der Waal et al. 2007).

There are a number of relevant caveats to this type of model. First, in its strict form it assumes that agents have the capacity to compute the costs and benefits of many different real-world policy packages, and therefore some level of cognitive understanding of the mechanisms at play. While the logic of a fixed benefit (transfer) and an income-dependent cost (tax) may be fairly intuitive to many people, it becomes much more complicated in a real-world setting with an immensely complex set of state institutions and a multitude of different types of income and taxation, where a large number of policy packages are pitted against each other, and where the relationship between the different policy proposals is much more opaque. Individuals who find it harder to engage in the rather complicated calculations necessary to evaluate and contrast the marginal effects of different policy packages, may therefore simply be less likely to optimize correctly (or at all). To further underscore this, there is a literature showing that the “quality” of economic decision making (i.e. consistency, compliance with monotonicity, proneness to fallacies) does in fact increase with cognitive performance (e.g. Burks et al. 2009, Choi et al. 2014, Oechssler et al. 2009).

Second, a long tradition in political science emphasizes the role of the so-called “impressionable years”: environmental influences on attitudes tend to be most important during late adolescence and early adulthood (Jennings and Niemi 1981, Alesina and Fuchs-Schündeln 2007, Giuliano and Spilimbergo 2014), after which they tend to remain fairly stable (Kustov et al. 2021, O’Grady 2019). This actualizes the role of class *background*: rather than simply probing contemporary and transient economic conditions, we ought to carefully consider the influence of early perceptions of relative affluence as captured by, for example, the economic standing of one’s parents. Further underscoring this, a recent study using a twin design found that actual wealth fails to exert a substantial causal influence on economic ideology, and that naive correlations between the two are instead mostly driven by factors in the family environment (Ahlskog and Brännlund 2021).

Following from this, I propose that rather than directly influencing economic policy preferences, cognitive performance (and hence genes associated with cognitive performance) influences the capacity to *correctly optimize* policy preferences, or, put differently, to correctly deduce one’s class interest. Furthermore, the perception of this class interest will to a large extent be shaped during the impressionable years by the relative economic standing one’s parents. Departing from traditional models in political economics, this leads to the following hypotheses:

**H1a:** Genetics associated with cognitive performance should be associated with *less* economic conservatism in individuals from *poor* socioeconomic backgrounds.
**H1b:** Genetics associated with cognitive performance should be associated with *more* economic conservatism in individuals from *affluent* socioeconomic backgrounds.

In summary, we should expect a GxE effect on economic ideology, where socioeconomic background moderates, and in fact reverses, the effect of genetics associated with cognitive performance. In the broader taxonomy of GxE effects, one can note that the focus here is on a genetic effect being moderated by the environment, as opposed to a genetic factor moderating a main environmental effect (though statistically equivalent, others have noted there are theoretical differences that can influence, for example, gene selection – e.g. Zhang and Belsky 2020). Furthermore, arguably the most important distinction is between situations where the magnitude but not the sign of a genetic effect may differ, and situations where even the sign may change (sign-discordance). These have previously been called “dimmer”-vs. “lens”-effects (Domingue et al. 2020). Since the proposed hypotheses imply a reversal of the sign of the genetic effect, this would indicate a lens-effect. This has crucial methodological implications discussed below.^2^

It is also important to delineate these hypotheses from previous arguments regarding the sign and mechanism of the effect of cognitive performance on economic ideology. To begin with, a case has been made that actual resources are a mediating factor: individuals with higher cognitive performance are assumed to have higher earnings potential, and therefore, on average, to be less likely to be net beneficiaries of taxation and redistribution (Jedinger and Burger 2022, Ganzach 2020). This argument is related in that it connects cognitive performance to an implicit cost-benefit framework. The difference lies in that it does not assume that cognitive performance is connected to rationality itself, but simply to the expected “payoff,” and therefore does not predict an interaction effect. It is also worth keeping in mind that possible resource effects of cognitive performance on political preferences require that either a) preferences actually change friction-free as a function of resources, in which case they must not be very sticky, or b) are formed during the impressionable years based on accurate expectations of future earnings informed by knowledge of one’s cognitive performance relative to others (“I’m smarter than my peers, therefore I will make more money than them in the future – when I do, I don’t want to pay for their welfare checks”). The plausibility of the latter mechanism can be debated.

Another argument is that cognitive performance disposes one toward right-leaning economic policies (“thinking like an economist”) regardless of personal circumstances because those policies are somehow “more logical” (Caplan and Miller 2010). Notwithstanding the debate that could be had around the premise of such an argument, it is worth pointing out that the mechanism I propose is, though in a different way, rather precisely aligned with this line of reasoning – in both cases, cognitive performance is presumed to make you more “rational.” The key difference lies in acknowledging that the more logical choice of economic policies will depend on the circumstances – such as socioeconomic background.

One can also distinguish the results from those stemming from extremism and context theory (Sidanius 1985). Extremism theory, on this account, predicts that cognitive performance should have a moderating effect, such that it makes people align more centre. If socioeconomic environment is a primary determinant of economic attitudes, extremism theory would lead to the opposite pattern of results: individuals in poor/rich environments will be leftists/rightists, but with higher cognitive performance they will move towards the centre. Meanwhile, context theory predicts the opposite: questioning the status quo (presumably the “centre”) and formulating independent viewpoints requires a certain amount of cognitive flexibility. Cognitive performance would then have *de*-moderating effects on ideology, which under certain circumstances could lead to the same pattern as the one I propose. More on how to test this can be found in the methods section.

A similar literature also exists on the relationship between education and ideology (Dunn 2011, Rasmussen et al. 2021, Weakliem 2002). It has been argued that since education confers resources, and scarcity of resources (level of inequality) varies between contexts, we should expect effects of education to also be contextually contingent. For example, one study finds that educational effects on economic conservatism are amplified in impoverished contexts (Rasmussen et al. 2021). Since cognitive performance can also arguably have downstream resource effects, the same logic could very well apply. This dynamic is related to but quite distinct from the hypotheses proposed here, however, in that the relative effects here are thought to be a function of relative affluence within a given context, rather than the degree of inequality or competition over resources in that context. One does not negate the other – although the focus here is not on contextual inequality, it may very well be that the degree to which the effect of cognitive performance differs between higher and lower socioeconomic background varies as a function of the local degree of inequality. Unfortunately this study would not be well-powered to detect a triple-interaction.

## Method and data

To credibly capture a causal GxE effect, one would ideally want a) random variation in the genetic factor of interest (to avoid the confounding effects of population stratification and genetic nurture), and b) an exogenous source of variation in the environmental moderator (to avoid any other confounders, including other genetic factors). Thanks to Mendel’s First Law, one way of obtaining random variation in the genetic factor is to move from the cross-sectional level of analysis to looking at differences between siblings. Fraternal (i.e. dizygotic, or DZ) twins are particularly helpful in this regard, since age/cohort effects are automatically partialled out as well (which could increase statistical precision over using siblings).

This study relies on a design leveraging within-family differences among DZ twins in economic conservatism and a polygenic index (PGI) of cognitive performance, interacted with between-family socioeconomic background (henceforth, family SEI). This provides a reasonable argument for causal identification of the effect of the PGI, but less so with the identification of family SEI, which may be confounded by a range of things (including genetics). However, family SEI does not suffer from reverse causation, and is not subject to self-selection effects (since twins cannot choose their parents). A number of precautions can be used to safeguard against other confounding in family SEI, such as controlling for all local-level confounding by using only within-parish variation, which I will return to below. The empirical model, in short, relies on differencing economic conservatism (Δ*Y_j_* = *Y*_1*j*_ – *Y*_2*j*_) and the PGI of cognitive performance (Δ*G_j_* = *G*_1*j*_ – *G*_2*j*_) within twin pair *j* and fitting the following equation:

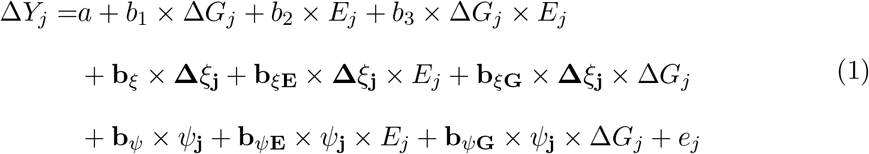

where *E* is family SEI, *ξ* is a vector of within-family controls and *ψ* is a vector of between-family controls. While this specification can be fit in a number of ways (including varying coefficients multi-level modeling), I have chosen what is arguably the most straight-forward and conservative approach, namely first transforming pairs of individual observations to single twin pair observations by differencing the within-family variables. In all results, observations therefore represent complete twin pairs as opposed to individuals.

The proposed hypotheses imply three things. First and foremost, the interaction coefficient *b*_3_ should be positive, such that an increase in E leads to a more positive effect of *G*. That is, the effect of the cognitive performance PGI should be more positive in families of higher SEI. However, since the hypotheses posit different *signs* of the genetic effect, this is not sufficient. We should also observe a negative marginal effect of *G* conditional on low family SEI, and a positive marginal effect of *G* conditional on high family SEI.

As stated above, using differences in a PGI within families has the benefit of having a credible causal interpretation. The point estimate, however, is going to be biased toward zero for two reasons. First, the PGI is a noisy measure of the true additive genetic factor (the precision is limited by the size of the GWAS discovery sample), meaning that it is affected by attenuation bias (Becker et al. 2021). Second, to the extent that genetic nurture effects are not perfectly correlated with the direct genetic effects, they essentially turn into *more* noise when differencing the PGI within twin pairs, thereby further exacerbating attenuation bias (Trejo and Domingue 2018). It is therefore important to keep in mind that effect size estimates are likely to be overly conservative.

The proposed empirical model can be distinguished from related but different approaches to using family data and a PGI to estimate GxE effects. One special case would be to use identical twins and within-family variation in an environmental exposure, but between-family variation in a PGI (since identical twins will also have identical PGIs). This would provide better identification of the environmental moderator. Due to aforementioned problems with population stratification and genetic nurture effects, it provides much poorer identification of the PGI effect (see e.g. Morris et al. 2020). Similarly, one could use dizygotic twin pairs and leverage within-pair variation in both the PGI and an environmental exposure. However, both of these designs preclude an environmental factor shared by twins, which is what we’re after in the present case. Thus, in this setting, we will need between-family variation in the environmental exposure (though with extensive controls) and live with the possibility of confounding for the moderator of interest.^3^

I argue that this is, nonetheless, a reasonable path forward. First, for most social scientists, it is likely going to be more important to credibly identify the genetic effect. While there is by now a substantial body of research documenting genetic influences on outcomes of interest for social scientists, genetic factors are still some distance away in many disciplines from being widely accepted as part-and-parcel of the toolkit for understanding variation in socially relevant human traits. If one contrasts identifying a credible “gene-by-something” interaction versus a “something-by-environment” interaction, the former should be of substantially larger interest. Second of all, we arguably have a better understanding of the possible confounders for something like family background than we do of the influences of genetic population stratification. We are therefore better placed to provide a nuanced interpretation of the results when genetic effects are clearly identified and environmental effects less so, than vice versa.

Something should also be said about the suitability of using a PGI derived from an additive GWAS in modeling this type of interaction. The character of the proposed interaction is what has been called a “lens”-effect, indicating that the effect of the genetic index will change signs depending on the environmental moderating factor (Domingue et al. 2020). This is in stark contrast to, for example, “dimmer”-effects, where the effect of the genetic index is simply thought to be stronger or weaker, but retain its sign. A PGI derived from an additive GWAS will capture the average treatment effect on the phenotype in question, and an interaction that is thought to span different signs may therefore have been completely washed out already at the discovery stage. In this regard, it is important to note that the PGI used in this study is *not* derived from a GWAS of economic conservatism, but of cognitive performance, and the moderating mechanism implied by the theory happens *after* the PGI target phenotype. This means we will be able to find a lens-effect despite the PGI being based on an additive GWAS of cognitive performance. In fact, the theory does rely on a consistent and monotonous effect of cognitive performance-linked genetics with the target *endo*phenotype (trait cognitive performance), after which the effect on economic ideology is moderated by the environment.

### Why use a genetic measure of cognitive performance?

A question that some may raise is why this should be approached as a GxE hypothesis at all, and not simply as a conventional interaction effect: i.e., why use a genetic measure of cognitive performance, as opposed to a more proximal trait-level measure (such as an IQ test)? Apart from the substantive argument that it is clearly of special interest to understand the precise etiology of previously established genetic effects on ideology, and what these genetic effects actually mean, there are, in my view, a number of purely methodological arguments that should be discussed. To start with, there are two possible arguments against the GxE route.

First, it is clear that a genetic measure will have larger measurement error than trait-level measures. Polygenic indices do not yet capture the full previously established genetic influence on cognitive performance, and genetics, furthermore, is not the only influence on the actual trait. This means that the amount of variation in real underlying individual cognitive ability that is captured is going to be smaller than if one were simply using an IQ test. Whereas the measurement error with an IQ test is just the measurement error of the instrument itself in capturing cognitive performance, the precision of a polygenic index is a function of the noise in the PGI *and* the heritability of the trait. This is the main drawback. The larger amount of measurement error, since it’s in the predictor, will also lead to larger attenuation bias, which in this case also negatively affects power to detect an interaction effect. As mentioned before, this will mean that any effects that are found are conservatively estimated.

Second, although as argued above, within-DZ pair differences in a genetic measure provide reasonable grounds for a causal interpretation, due to pleiotropy the causal mechanism need not in fact be exclusively through cognitive capacity, to the extent that cognitive capacity is *genetically correlated* with some other causal factor. While the difference in the genetic measure of cognitive performance between two fraternal twins is therefore plausibly causally related to other downstream differences (i.e. in ideology), it is not *necessarily* just an effect of actual trait cognitive performance. This does not detract from the causal interpretation, but may render conclusions about the proposed mechanism less definitive. It is important to add, however, that in a different sense this would be true (though perhaps to a smaller extent) with a trait-level measure as well, since the trait-level measure is also going to be correlated with other factors.

There are, however, even stronger arguments in favor of using a genetic measure. The first two arguments in favor simply regard the fundamental issue of data availability – both in general and in the specific sample at hand – and the final one regards implications for the identification strategy. First, if one has a genotyped sample, no additional data on cognitive capacity is needed – the required polygenic index (or any other polygenic index, for that matter) can be computed from just the genetic data. Additionally, no other concerns regarding, for example, possible discrepancies between different types of survey instruments used to measure cognitive performance apply.

Second, the proposed empirical strategy relies on complete twin pairs, and available IQ tests (in the case at hand) come from military conscription data. The latter fact means that IQ data is available almost exclusively for male subjects. When complete dizygotic pairs are required, this means that not only the female-female pairs are lost, but also both data points in the male-female pairs. If one were to, instead, settle for a discordant MZ twin design with trait-level measures, one has to contend with the fact that there are far fewer (about half as many) identical twins than fraternal twins, and that only half of the pairs would be male-male pairs. In sum, the resulting total loss would be a minimum of 3/4 of a dizygotic sample and 1/2 of a (much smaller) monozygotic sample. Add to this the obvious issue of external validity if results are based on males exclusively.

Third, and more importantly, measurement error in an actual IQ test can be correlated with other factors that influence the outcome – perhaps especially so on a test performed during military conscription – thus introducing bias. This becomes especially salient in light of the results from Burger et al. (2020), that part of the observed relationship between cognitive performance and political ideology is driven merely by the fact that individuals high in right-wing authoritarianism have lower motivation to *perform* well on cognitive tasks in the first place. This is not the case with a genetic measure, which is assigned at conception and subsequently unaltered throughout the life cycle. Another way of viewing this is to say that within-pair variation in IQ test results is indeed going to be purged of all confounders that are shared among siblings – but not all confounders. Importantly, half of the genetic confounding, and all confounders residing in the unique environment remain – consider, for example, if the siblings have different groups of friends, which differ in their socializing effects on both intellectual development and political ideology. On the contrary, variation in a genetic measure between dizygotic twins is – as repeatedly stated already – random, meaning that these confounders are ruled out by design.

### Data

I make use of a sample of DZ twin pairs from the Swedish Twin Registry (Zagai et al. 2019). The sample contains all complete genotyped DZ twin pairs that have answered the relevant items of the SALTY survey, which was conducted in 2009/2010 among twins born between 1943 and 1958. The twins have also been connected to rich registry sources for things like education and income, as well as to the intergenerational registry, allowing the addition of the same register variables for the parents of the twins. A separate population-based register dataset has been used to obtain information about context/neighborhood SEI. All variables used are described below.

### Economic ideology

The main outcome is economic ideology. The SALTY survey contains a battery of political preference questions, where the respondent is asked to indicate, on a five-point scale ranging from “Strongly disagree” to “Strongly agree” what their opinion is about 34 different policy proposals. The questions span a large range of different types of preferences, from welfare policy to environmental and foreign policy. I have used these to construct an operational measure of economic conservatism.

The outcome variable is constructed in three steps, to get as precise a measure as possible. First, I identified the items in the survey that have strong face validity as measures of economic ideology, involving regulation, taxation, redistribution and financing of the public sector. The twelve items I included are as follows:

- Decrease the public sector
- Lower welfare payments
- Lower taxes
- Sell publically owned companies to private buyers
- Run more healthcare in the private sector
- Promote more free trade in the world
- More private schools^4^
- More freedom for companies
- Decrease the influence of the financial market over politics
- Keep property taxes
- Keep maximum fees in public child care
- Decrease economic inequality

These items have a Cronbach’s alpha of 0.75. The second step was to fit a structural equation model for all identified items, to gauge the relative importance of each item for economic ideology. To avoid overfitting, this was done out of sample using one individual from each monozygotic twin pair (since the main sample only uses dizygotic twin pairs). The RMSEA for all items out of sample was .077, which should be considered a satisfactory fit. The last step was to construct the outcome as a mean of all the included variables, weighted by the coefficients estimated for the latent factor, and rescaling the measure to the original five-point range (rescaling is necessary for the calculation of ideological constraint below).

### Ideological extremism

To rule out that possible interaction effects are artefacts of simple moderation/de-moderation effects, in line with extremism or context theory (i.e. cognitive performance pushing people either to the centre or towards the edges of the scale), models will also be tested with measures of extremism defined as the absolute distance from the scale midpoint.

### Ideological constraint

To control for the degree of constraint or coherence of an individual’s economic ideology, I use the method proposed by Barton and Parsons (1977) by calculating the standard deviation of the included items from their weighted mean (as defined above), per respondent. The intuition behind this is fairly straight forward: a completely ideologically consistent individual will answer symmetrically on all items (i.e. a fully economically conservative person will answer “Completely agree” on all items positively related to economic conservatism, and “Completely disagree” on all items negatively related to economic conservatism, and vice versa) and therefore have a very low standard deviation, while an ideologically inconsistent individual will tend to answer more randomly, and therefore have a higher standard deviation.

### Family SEI

Family SEI is measured using data on parental education and income, for both the mother and the father. To partial out all possible local or regional geographic confounders, I used a separate full-population register dataset to calculate the parish- and sex-specific percentiles^5^ for education and income for the two parents (among all individuals in the mother’s parish having reached the age of 20), after regressing out parental birth year fixed effects to account for age related earnings- and education profiles. These four percentile ranks were then averaged and standardized by birth cohort of the twins to get the final measure of family SEI. Due to data limitations, this SEI measure is based on census data for the year 1970.^6^ Since the SALTY sample were born between 1943 and 1958, the genotyped respondents will have been between 12 and 27 years of age in 1970. Some portion of them will thus have left their parental home at the time of measurement.

### PGI of cognitive performance

The genetic measure used in this study is a polygenic index of cognitive performance (henceforth, CogPerf PGI), taken from the recently established PGI repository (Becker et al. 2021). To maximize the explanatory power of the cognitive performance PGI, I have opted to use the multi-trait (MTAG) PGI, which is constructed with educational attainment as its supplementary phenotype. The expected explanatory power (as measured by the incremental *R*^2^) of this PGI for trait cognitive performance is 15.77% (Becker et al. 2021). Due to possible genetic nurture effects and residual population stratification, this is going to be substantially smaller when differencing between dizygotic twins. The predictive power of the PGI in within-family models for both a measure of IQ (based on military conscription tests for males) and education years is presented in table 1. These results show that a) the PGI significantly predicts both IQ test results (*t* = 6.71) and education years (*t* = 8.71), b) the effect is roughly twice as large for IQ, and c) the *R*^2^ for the target phenotype is 5.1% even within DZ twin pairs, i.e. about a third of the expected between-family *R*^2^.

**Table 1:**
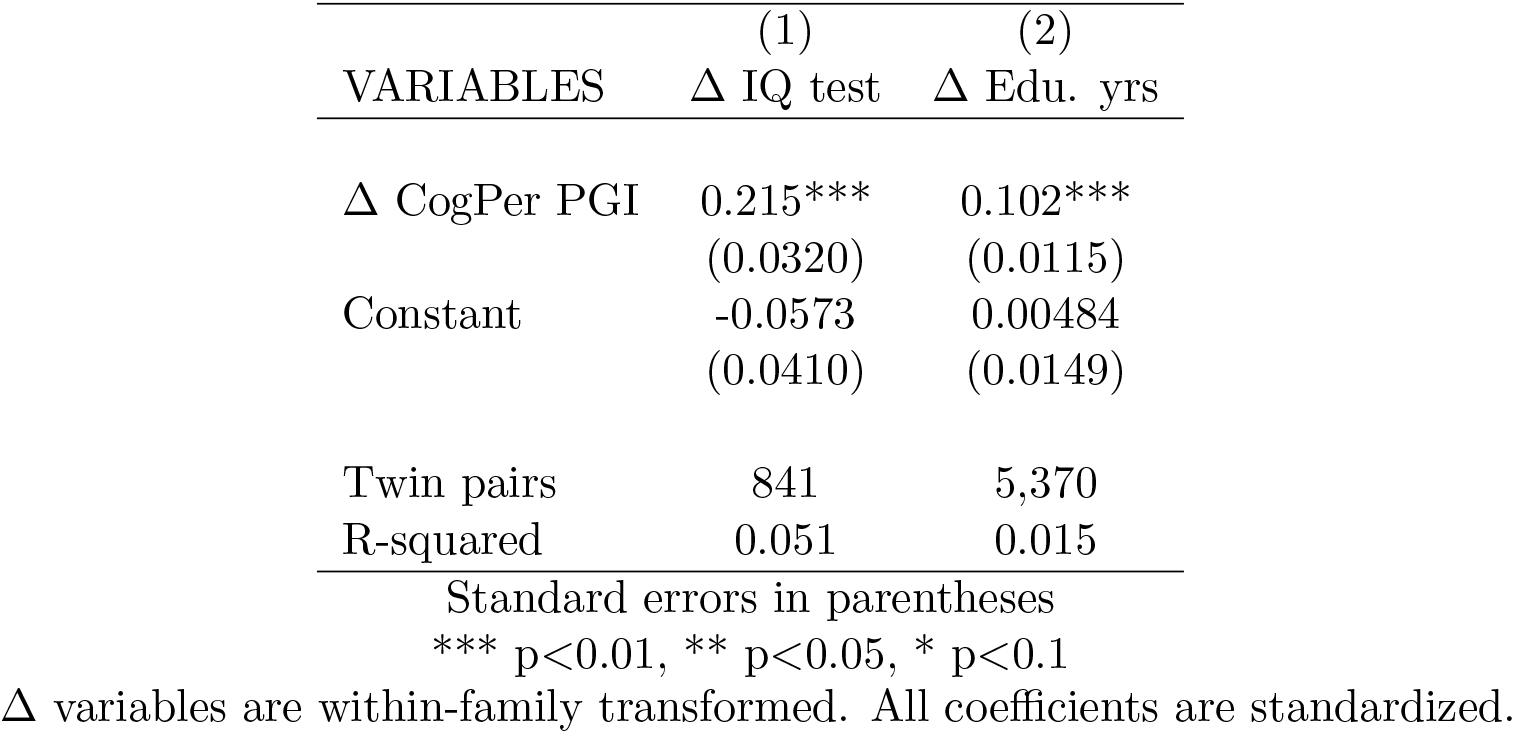
Within-pair prediction

Also note that since within-family models are used, genomic principal components (traditionally used to capture some of the confounding effects of population stratification) are left out.

### Controls

Apart from the sex of the twins (included in all models as a within-family control), a number of covariates are introduced to safeguard against confounding of variation in family SEI, to increase precision, and to exclude alternative explanations.

First and foremost, intergenerational transmission makes it important to distinguish the familial socioeconomic background from the current economic situation of the respondent. To do this, I constructed a similar measure based on education and income of the individual respondent at the time of the survey, based on the LISA register database from Statistics Sweden. Education years was measured in 2009, i.e. at the time of the survey. Income, further, was measured as the average income in the ten years preceding the survey to minimize random variability. Birth year fixed effects were also regressed out to account for life-cycle effects on income. Again, percentiles of education and income were taken and the two percentiles were averaged with equal weight. This measure is included as a within-family control in a separate model to see the level of mediation by current SEI.

Second, I also control for the between-family average SEI as well as population size of the childhood parish. The parish SEI is computed by taking the sex-specific mean income and education years per parish, taking percentiles of each, and averaging these four percentiles.

Lastly, to decrease the risk of mistakenly identifying a gene-gene interaction or a non-linear main genetic effect (such that effects at either end of the SEI spectrum somehow reflects average differences in the PGI), I also include between-family average PGI’s and average PGI’s squared for both the cognitive performance PGI as well as all other available indices of behavioral phenotypes with an expected single-trait incremental *R*^2^ of at least 2%, excluding health traits, in the PGI repository (Becker et al. 2021). This includes educational attainment, chronotype, risk seeking, adventurousness, extraversion, neuroticism and subjective wellbeing (indices for math ability were excluded due to their high colinearity with the index for cognitive performance). I also included height, since it is currently the most predictive PGI in the repository and height is known to be a) genetically correlated with cognitive performance (Keller et al. 2014), and b) phenotypically correlated with political preferences (Arunachalam and Watson 2018).

As detailed in Eq. 1, in all models with controls included, their respective interactions with the two main variables of interest (the within-pair difference in the cognitive performance PGI, and the measure of family SEI) are also included (Keller 2014).

## Results

The main results are shown in Table 3. We can see in models 1 and 2 that the average effect of the cognitive performance PGI is not distinguishable from zero. From the perspective of previous hypotheses on the connection between cognitive performance and economic ideology, we would have stopped at this point and added a null result to the literature.

**Table 2:**
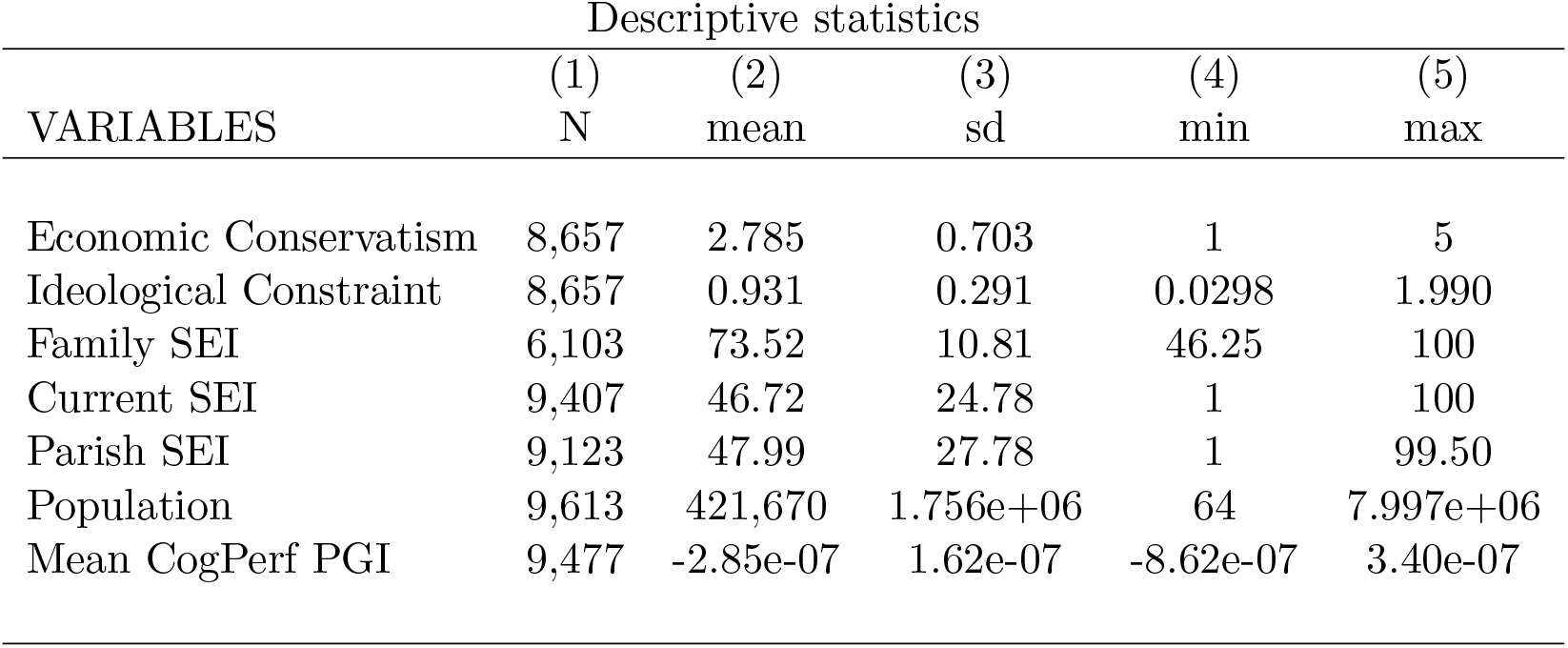
Descriptive statistics

| VARIABLES | Descriptive statistics |  |  |  |  |
| --- | --- | --- | --- | --- | --- |
|  | (1)<br>N | (2)<br>mean | (3)<br>sd | (4)<br>min | (5)<br>max |
| Economic Conservatism | 8,657 | 2.785 | 0.703 | 1 | 5 |
| Ideological Constraint | 8,657 | 0.931 | 0.291 | 0.0298 | 1.990 |
| Family SEI | 6,103 | 73.52 | 10.81 | 46.25 | 100 |
| Current SEI | 9,407 | 46.72 | 24.78 | 1 | 100 |
| Parish SEI | 9,123 | 47.99 | 27.78 | 1 | 99.50 |
| Population | 9,613 | 421,670 | 1.756e+06 | 64 | 7.997e+06 |
| Mean CogPerf PGI | 9,477 | -2.85e-07 | 1.62e-07 | -8.62e-07 | 3.40e-07 |

**Table 3:** Main results, Economic Ideology

| VARIABLES | (1)<br>$\Delta$ EC | (2)<br>$\Delta$ EC | (3)<br>$\Delta$ EC | (4)<br>$\Delta$ EC | (5)<br>$\Delta$ EC | (6)<br>$\Delta$ EC |
| --- | --- | --- | --- | --- | --- | --- |
| $\Delta$ CogPerf PGI | -0.0182<br>(0.0275) | -0.0160<br>(0.0276) | -0.00352<br>(0.0470) | -0.134<br>(0.159) | -0.198<br>(0.162) | -0.145<br>(0.159) |
| Family SEI |  |  | -0.000973<br>(0.0601) | 0.176<br>(0.209) | 0.165<br>(0.213) | 0.171<br>(0.209) |
| $\Delta$ CogPerf PGI × Family SEI | | | 0.0741***<br>(0.0282) | 0.0698**<br>(0.0309) | 0.0749**<br>(0.0319) | 0.0661**<br>(0.0310) |
| $\Delta$ Current SEI | | | | | 0.0460<br>(0.0383) | |
| $\Delta$ CogPerf PGI × $\Delta$ Current SEI | | | | | 0.0454<br>(0.0287) | |
| $\Delta$ Ideological Constraint | | | | | | -0.0217<br>(0.0382) |
| $\Delta$ CogPer PGI × $\Delta$ Ideol Const | | | | | | -0.0401<br>(0.0284) |
| Constant | -0.0346<br>(0.0365) | -0.0519<br>(0.0623) | -0.0521<br>(0.0625) | -0.229<br>(0.208) | -0.244<br>(0.211) | -0.252<br>(0.209) |
| Twin pairs | 857 | 857 | 857 | 857 | 819 | 857 |
| R-squared | 0.001 | 0.002 | 0.014 | 0.106 | 0.115 | 0.109 |
| Controls | NO | YES | YES | YES | YES | YES |
| Mean/Mean<sup>2</sup> PGIs | NO | NO | NO | YES | YES | YES |

Standard errors in parentheses
\*\*\* $p < 0.01$ , \*\* $p < 0.05$ , \* $p < 0.1$
$\Delta$ variables are within-family transformed. Controls are for sex, parish SEI and parish population size and their interactions with $\Delta$ CogPerf PGI and Family SEI. Mean/Mean<sup>2</sup> PGIs are the full set of family average PGI's and family average PGI's squared as well as their interactions with $\Delta$ CogPerf PGI and Family SEI. Model 4 contains an interaction term between Family SEI and $\Delta$ Current SEI. Model 5 contains an interaction term between Family SEI and $\Delta$ Ideological Constraint. All coefficients are standardized.

However, the hypotheses posited in this paper requires us to investigate the moderating effect of family SEI. Moving to model 3, I therefore interact the within-pair differences in the cognitive performance PGI with the cross-pair differences in family SEI. We now see that the average null effect does in fact hide heterogeneity: without the mean PGI controls, the coefficient for the interaction is positive and significant (*p* = .009), indicating that the effect of the cognitive performance PGI on economic conservatism increases by the level of family SEI. With the full set of mean and squared family PGIs in model 4, this result remains (*p* = .024) and is only marginally weakened, indicating that the identified interaction is not likely to be driven by a gene-gene interaction with any of the included genetic indices.

Moving to the marginal effects plot of model 3 in Figure 2, we can further inspect the character of this interaction. The differences are striking: at the lower levels of relative SEI, the effect of the cognitive performance PGI on economic conservatism is negative, meaning that a higher PGI in a poor family leads to more attitudes in favor of taxation and redistribution. Meanwhile, at the higher levels of relative SEI, the effect is instead positive, meaning that a higher PGI in a rich family leads to more attitudes in favor of cutting welfare spending and relying more on markets. The crossover point is around zero. The effect sizes at either end of the family SEI distribution is small to moderate, corresponding to roughly at most .2 standard deviations in economic ideology per standard deviation in the cognitive performance PGI. These effect sizes are unadjusted for measurement error and are therefore most likely substantially larger in reality.

**Figure 1:**
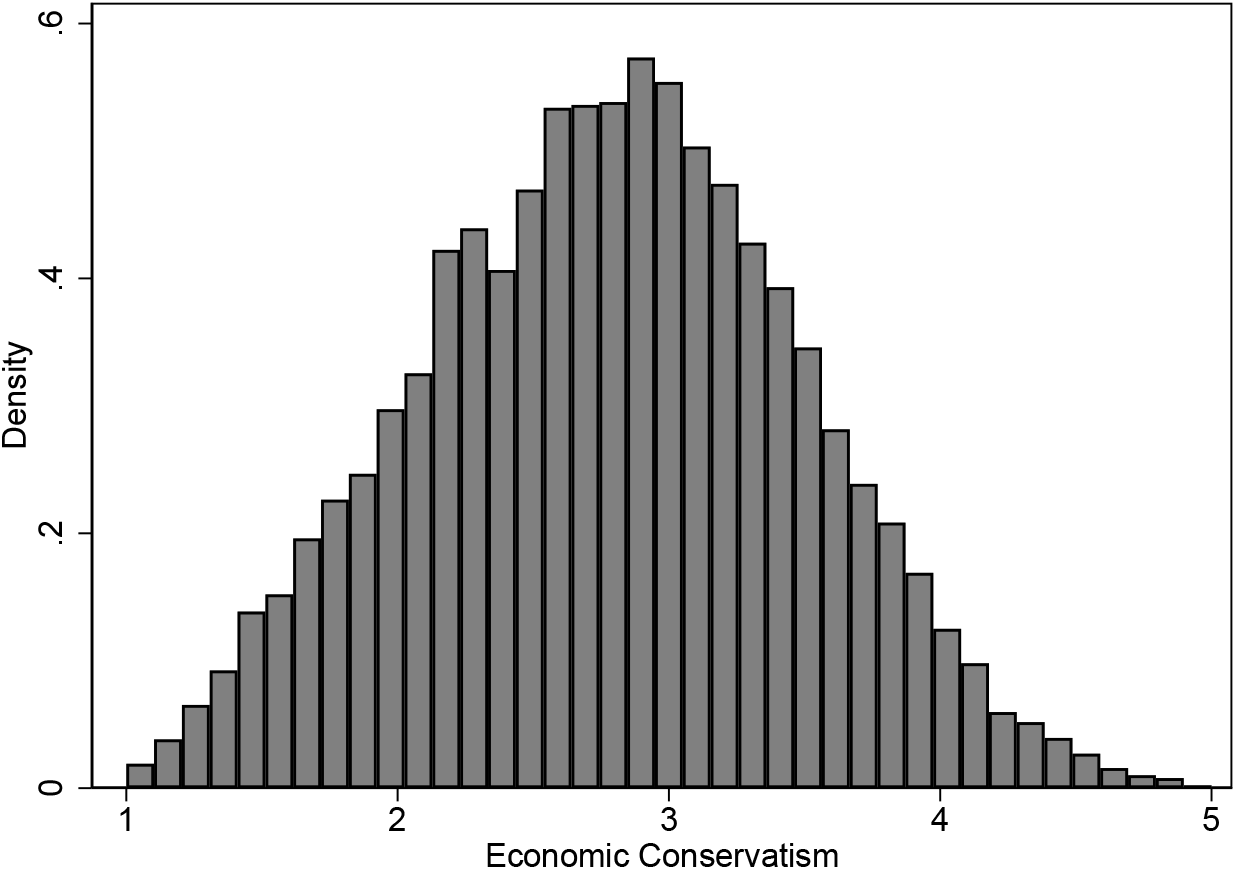
Distribution of economic conservatism

**Figure 2:**
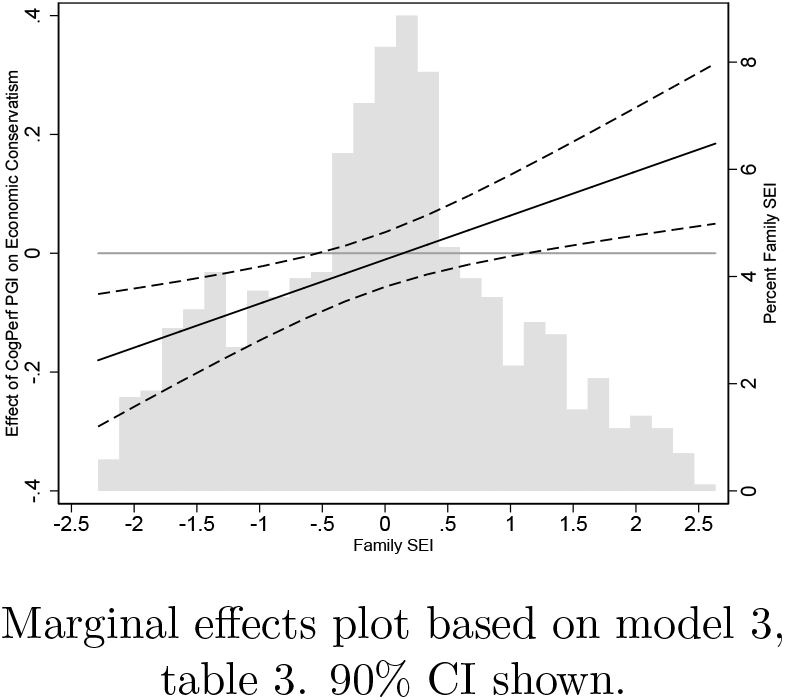
Marginal effects plot, CogPerf PGI effect on Economic Conservatism

An alternative explanation for this result is that family SEI is merely a proxy for economic conditions later in life. To investigate to what extent this is true, model 5 in Table 3 also includes within-pair differences in a measure of current SEI (as well as interactions between this measure and both family SEI and the PGI). As indicated by the lack of a significant interaction term between the cognitive performance PGI and current SEI, and the fact that the original interaction term increases somewhat in magnitude, there is no evidence for this alternative explanation. This speaks against a resource interpretation of the cognitive performance effects: the effect of the PGI does not in fact appear to be mediated (or moderated) by resources later in life.

Another possibility is that the estimated interaction is an artefact of an effect on increased ideological constraint. Since the economic ideology scale has defined endpoints, it is conceivable that the environment (as measured by family SEI) moves one to the left or to the right, and cognitive performance merely makes one’s ideological position more consistent and therefore less “biased” towards the middle.^7^ To safeguard against this, model 6 adds the measure of ideological constraint for economic ideology (as well as interactions between this measure and both family SEI and the PGI). Again, we see that there appears to be no effect of the constraint measure, and that the interaction between family SEI and the cognitive performance PGI remains at roughly the same point estimate.

It is also possible that the interaction effect itself is non-linear, and that the results above are driven by just one rather than both ends of the SEI spectrum. One way of investigating this is to allow for full flexibility in the functional form of the interaction by performing rolling regressions with limited samples across family SEI. Figure 3 shows rolling regressions with a window size of ±1 unit of family SEI. We can see that there is some fluctuation across the distribution of family SEI, but that the there is no hint of a U-shaped relationship. More importantly, the predicted pattern is still very clear: negative effects on the low end and positive effects on the high end. The curve is fairly monotonous, though not perfectly linear, but the deviation from linearity appears small enough that a linear interaction term provides a reasonable approximation.

**Figure 3:**
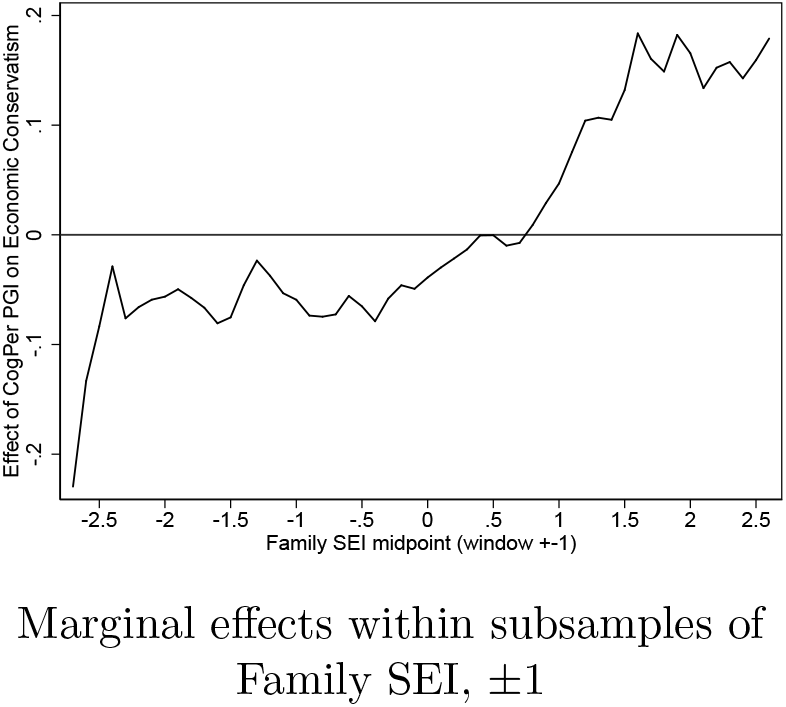
Rolling regressions, CogPerf PGI effect on Economic Conservatism

To further distinguish the identified interaction from de-moderation effects, such as those proposed by context theory (Sidanius 1985), we can look at the main additive effect of the cognitive performance PGI on “extremism,” i.e. the distance from the scale midpoint. These results can be found in table 4. The point estimates show a *moderation* effect (a higher cognitive performance PGI leads to less extremism), indicating that the main results are not driven by de-moderation effects.

**Table 4:** Extremism.

| VARIABLES | (1)<br>$\Delta$ EXT EC | (2)<br>$\Delta$ EXT EC |
| --- | --- | --- |
| $\Delta$ CogPerf PGI | -0.0210<br>(0.0220) | -0.0266<br>(0.0227) |
| $\Delta$ Ideological Constraint | | -0.0757***<br>(0.0289) |
| $\Delta$ Current SEI | | 0.00380<br>(0.0281) |
| $\Delta$ Sex | -0.0384<br>(0.0420) | -0.00595<br>(0.0442) |
| Constant | 0.00737<br>(0.0292) | 0.0179<br>(0.0298) |
| Observations | 1,337 | 1,273 |
| R-squared | 0.001 | 0.006 |

Standard errors in parentheses
\*\*\* $p < 0.01$ , \*\* $p < 0.05$ , \* $p < 0.1$ $\Delta$ variables are within-family transformed.

In summary, there appears to be good evidence of an interaction effect between family SEI and a cognitive performance PGI on economic conservatism. This interaction is such that the effect of the PGI is negative among individuals coming from less affluent families, but positive among individuals coming from more affluent families.

## Discussion

In this paper I have connected to previous studies arguing that some portion of genetic effects on ideology originate in genetic factors connected to cognitive performance. However, the proposed mechanism does not involve an effect on policy preferences per se, but on to what extent people are able to optimize their policy preferences. Since rational preference formation over complex policy packages in a modern interconnected society require some degree of cognitive computational complexity, we should expect more cognitively “sophisticated” individuals to be more likely to be able to rationally optimize their policy preferences. Moreover, if it is true that political preferences are “sticky,” this implies that said optimization process must have been performed, at least to a large extent, against the backdrop of the economic conditions prevalent during one’s impressionable years.

The first thing to note about the results is that the main, unmoderated effect of a cognitive performance PGI on economic conservatism in this sample of Swedish respondents, is zero. That is, on average, having more alleles associated with cognitive performance (and more specifically, having more alleles than your twin) did not dispose these respondents to becoming either more favorable to taxation and redistribution or more economically conservative. However, this lack of average effect did, in line with the proposed hypotheses, hide substantial underlying heterogeneity – not just in the sense that it was stronger than zero among some people, but that the effect was positive among some and negative among others. This differential, along socioeconomic lines, has a strong theoretical rationale: economic ideology is arguably not a one-size-fits-all phenomenon, where it is always “smarter” to prefer one set of policies over another. On the contrary, by the relational nature of these preferences, rational economic policy preference formation can lead to diametrically different positions depending on the social class of the agent.

In absolute terms, the size of the effect at the margins of the SEI distribution is small to moderate. It should be noted, however, that both the effect sizes and the model R^2^ are likely to be fairly drastically underestimated. To begin with, the PGI is affected by a large amount of attenuation bias. Whereas the expected R^2^ of the used PGI is 15.77% (Becker et al. 2021), SNP heritability estimates (between families) of cognitive performance are in the range of 30-50% (Hill et al. 2018, Davies et al. 2016).^8^ This means that the PGI absorbs just around a third to half of the true additive heritability of the target endophenotype (if this proportion remains roughly the same when population stratification and genetic nurture is partialled out, the estimated coefficients in the results above should also capture about a third to half of the true effect size). Furthermore, if genetic nurture effects are present, the proportion of the PGI that captures genetic nurture will partially turn into more measurement error in within-family models and therefore exacerbate attenuation bias further (Trejo and Domingue 2018).

The main methodological perk of the design chosen in this study is the ability to credibly identify the causal influence of the genetic factor. However, this may come somewhat at the expense of the identification of the environmental moderator. In that sense, what has actually been identified should, strictly speaking, perhaps be labeled a “gene-by-something” interaction. That is, I can’t claim that the apparent moderating effect of family background on the main effect of the genetics of cognitive performance is, in fact, a causally moderating effect of family background, as opposed to something else merely correlated with family background. Several controls and robustness checks have been laid out to safeguard against the most salient obstacles to such an interpretation, but some will surely remain. One prospect is that the captured moderating effect actually represents a gene-by-gene interaction as opposed to a gene-by-environment ditto. This risk is substantially dampened by the inclusion of a wide range of family-level genetic factors *other* than for cognitive performance, but cannot be completely ruled out.

A final consideration is that the sample at hand is also unique in several ways, not only because they are twins. Two characteristics of this sample may impact their susceptibility to the proposed effects. The first is the country setting. The 20:th century political history of Sweden was overwhelmingly marked by the emergence of a strong labor movement, social democracy and the welfare state. Thus, the political context is one where the focal point of public political debate has tended to be on the precise economic ideological issues investigated here. Second, this is further accentuated by the age of the sample: having been born between 1943–1958 arguably puts the individuals square in the middle of the apex of the welfare state during their impressionable years. Yet, it is not clear whether this would be favorable or unfavorable conditions for the proposed hypotheses. On the one hand, we can confidently say that class-based mass politics is more salient in this context than elsewhere, which may serve as an environmental pressure on how people generally reason about politics. Thus, the theoretical premise that people optimize policy preferences based on something akin to class interest is more likely to be true here. On the other hand, previous research has shown that in the context of political participation, strong environmental cues may serve to dampen genetic influences, and vice versa (Oskarsson et al. 2022). It is therefore also possible that the strong environmental cues in this context makes the PGI effects themselves smaller.

The wider relevance of this work for other researchers using polygenic indices to predict various social and behavioral phenotypes also becomes clear once we consider that lens-type GxE effects will have a tendency to strongly bias average effects towards zero. An important implication of this is that a linear null result may not be a true null result, if the researcher has failed to consider the possible contingency of the sign of the effect on third variables. Moreover, using samples that are strongly self-selected on, for example, socioeconomic variables may also lead to worse external validity than usually expected if even the sign of the effect may differ in groups that are underrepresented among the research participants.

Finding out which specific gene-phenotype relationships are likely to be affected by this will ultimately have to be a theoretically grounded exercise. In this study, a moderating factor was proposed, based on existing models of preference formation, to be socioeconomic background. In other cases it might be something different altogether.

## Footnotes

1 As a salient example: in a large review of political preference formation from 2000, the possibility of genetic influences are not mentioned once (Druckman and Lupia 2000).

2 Additionally, there are taxonomies relating to the precise nature of the moderating effect of the environment. For example, Shanahan and Hofer suggest four different models (triggering, compensation, social control and enhancement) that serve to illuminate when a genetic effect is activated or not, or what the strength of the genetic effect is (Shanahan and Hofer 2005). Neither of the proposed models are applicable to lens-effects, however.

3 Though the empirical section of this paper also tests models with current SEI as opposed to family SEI, which can indeed be distinguished within pairs.

4 The specific type of private schools indicated in the item are so-called “freeschools,” i.e. privately run schools included in a voucher system.

5 Doing this per parish is essentially analogous to regressing out parish fixed effects.

6 Census data (FoB) is available at five year intervals from 1960–1990, but individual data for education and income only has partial coverage and is available in 1970.

7 To see why this is the case, consider an individual whose “true” ideological position is at the lowest possible level of economic conservatism. The only way to decrease ideological constraint from this point is to also increase the average level of economic conservatism. Therefore, at the extremes, we will only find individuals with very high ideological constraint.

8 The size depends on what cognitive domain one looks at, and also if one takes rare variants into account (Hill et al. 2018).

## References

Ahlskog, R. and Brännlund A. (2021). “Uncovering the Source of Patrimonial Voting: Evidence from Swedish Twin Pairs.” Political Behavior 44.

Alesina, A. and Fuchs-Schündeln, N. (2007). “Goodbye Lenin (or not?): The effect of communism on people’s preferences.” American Economic Review, 97(4).

Alford, J., Funk, C., Hibbing, J. (2005). “Are Political Orientations Genetically Transmitted?” APSR 99:153–167.

Anderson, D. and Davidson, P. (1943). Ballots and the Democratic Class Struggle: a study in the background of political education. Stanford, CA: Stanford University Press.

Arunachalam R. and Watson, S. (2018). “Height, Income and Voting.” British Journal of Political Science, 48(4).

Barton, AH. and Parsons, RW. (1977). “Measuring Belief System Structure.” Public Opinion Quarterly, 41(2).

Becker, J., CAP. Burik, G. Goldman, N. Wang … A. Okbay (2021). “Resource profile and user guide of the Polygenic Index Repository.” Nature Human Behavior, 5.

Bowles, S. and Gintis, H. (2011). A Cooperative Species: Human Reciprocity and its Evolution, Princeton University Press.

Burger, AM., Pfattheicher, S., Jauch, M. (2020). “The role of motivation in the association of political ideology with cognitive performance.” Cognition, 195.

Burks, SV., Carpenter, JP., Goette, L., Rustichini, A. (2009). “Cognitive skills affect economic preferences, strategic behavior, and job attachment.” PNAS, 19.

Caplan, B. and Miller, SC. (2010). Intelligence makes people think like economists: Evidence from the General Social Survey. Intelligence 6.

Carl, N. (2015). “Cognitive ability and political beliefs in the United States.” Personality and Individual Differences, 83.

Choi, S., Kariv, S., Müller, W., Silverman, D. (2014). “Who is (more) rational?” American Economic Review, 104(6).

Choma, BL., Sumantry, D., Hanoch, Y. (2019). “Right-wing ideology and numeracy: A perception of greater ability, but poorer performance.” Judgment and Decision Making, 114(4).

Clark, TN. and Lipset, SM. (1991). “Are Social Classes Dying?” International Sociology, 6(4).

Davies, G., Marioni, RE., Liewald, DC. …Deary, IJ. (2016) “Genome-wide association study of cognitive functions and educational attainment in UK Biobank (N=112 151).” Molecular Psychiatry, 21.

Dawes, CT., Cesarini, D., Fowler, JH., Johannesson, M., Magnusson PKE. and Oskarsson, S. (2014). “The Relationship between Genes, Psychological Traits, and Political Participation.” AJPS 58(4).

Deary, IJ., Batty, GD. and Gale, CR. (2008). “Bright Children Become Enlightened Adults.” Psychological Science 19(1).

Dick, DM., Agrawal, A., Keller, MC., Adkins, A., Aliev, F., Monroe, S., Hewitt, JK., Kendler, KS., Sher, KJ. (2015). “Candidate Gene-Environment Research: Reflections and Recommendations.” Perspect Psychol Sci, 10(1).

Domingue, BW., Trejo, S., Armstrong-Carter, E., Tucker-Drob, EM. (2020). “Interactions between Polygenic Scores and Environments: Methodological and Conceptual Challenges.” Sociological Science, 7.

Druckman, JN. and Lupia, A. (2000). “Preference Formation.” Annu. Rev. Polit. Sci. 3:1–24.

Dunn, K. (2011). “Left-Right Identification and Education in Europe: A Contingent Relationship.” Comparative European Politics, 9:292–316.

Eidelman, S., Crandall, CS., Goodman, JA. and Blanchar, JC. (2012). “Low-Effort Thought Promotes Political Conservatism.” Personality and Social Psychology Bulletin, 38(6).

Scott Eidelman, Christian S. Crandall, Jeffrey A. Goodman, and John C. Blanchar-1View all authors and affiliations

Evans, G. (2000). “The Continued Significance of Class Voting.” Annu. Rev. Polit. Sci., 3:401–17.

Evans, G. and Tilley, J. (2012). “The Depolitization of Inequality and Redistribution: Explaining the Decline in Class Voting.” Journal of Politics, 74(2).

Feldman S. and Johnston, C. (2014). “Understanding the Determinants of Political Ideology: Implications of Structural Complexity.” Political Psychology, 35(3).

Ganzach, Y. (2020). “From intelligence to ideology: socioeconomic paths.” Personality and Individual Differences, 164.

Giuliano, P., and Spilimbergo, A. (2014). “Growing up in a recession.” The Review of Economic Studies, 81:787–817.

Hatemi, P. et al. (2014). “Genetic Influences on Political Ideologies: Twin Analyses of 19 Measures of Political Ideologies from Five Democracies and Genome-Wide Findings from Three Populations.” Behavior Genetics.

Hill, WD., Arslan RC., Xia, C. … Penke, L. (2018) “Genomic analysis of family data reveals additional genetic effects on intelligence and personality.” Molecular Psychiatry, 23.

Jedinger, A. and Burger, AM. (2022). “Do Smart People Have More Conservative Economic Attitudes? Assessing the Relationship Between Cognitive Ability and Economic Ideology.” Personality and Social Psychology Bulletin, pp. 1–18.

Jennings, MK. and Niemi, RG. (1981). Generations and politics: A panel study. Princeton: Princeton University Press.

Keller, M., Garver-Apgar, CE., Wright, MJ., Martin, NG., Corley, RP., Stallings, MC., Hewitt, JK., Zietsch, BP. (2014). “The Genetic Correlation between Height and IQ: Shared Genes or Assortative Mating?” PLOS Genetics, 10(3).

Keller, M. (2014). “Gene x Environment interactions have not properly controlled for potential confounders: The problem and the (simple) solution.” Biological Psychiatry, 75(1).

Kustov, A., Laaker, D., Reller, C. (2021). “The Stability of Immigration Attitudes: Evidence and Implications.” The Journal of Politics, 83(4).

Ludeke, SG. and Rasmussen, SHR. (2018). “Different political systems suppress or facilitate the impact of intelligence on how you vote: A comparison of the U.S. and Denmark.” Intelligence, 70.

Manza, J., Hout, M. and Brooks, C. (1995). “Class Voting in Capitalist Democracies Since World War II: Dealignment, Realignment, or Trendless Fluctuation?” Annual Review of Sociology, 21:137–162.

Marx, K. (1887). Capital, vol I.

Meltzer, AH. and Richard, SF. (1981). “A Rationality Theory of the Size of Government.” Journal of Political Economy, 89(5).

Morris, TT., Davies, NM., Hemani, G. and Smith, GD. (2020). “Population phenomena inflate genetic associations of complex social traits.” Science Advances, 6(16).

Oechssler, J., Roider, A., Schmitz, PW. (2009). “Cognitive ability and behavioral biases.” Journal of Economic Behavior and Organization, 72(1).

O’Grady, T. (2019). “How do Economic Circumstances Determine Preferences? Evidence from Long-run Panel Data.” The British Journal of Political Science, 49(4).

Oskarsson, S., Ahlskog, R., Dawes, CT. and Lindgren, KO. (2022). “Persistent Inequalities: The Origins of Intergenerational Associations in Voter Turnout.” Journal of Politics 84(3).

Oskarsson, S., Cesarini, D., Dawes, CT., Fowler, JH., Johannesson, M., Magnusson, PKE. and Teorell, J. (2014). “Linking Genes and Political Orientations: Testing the Cognitive Ability as Mediator Hypothesis.” Political Psychology 36(6).

Oskarsson, S., Dawes, CT., Johannesson, M. and Magnusson, PKE. (2012). “The Genetic Origins of the Relationship between Psychological Traits and Social Trust.” Twin Research and Human Genetics 15(1).

Pereira, RD., Biroli, P., Galama, T., von Hinke, S., van Kippersluis, H., Rietveld, CA., Thom, K. (2022). “Gene-Environment Interplay in the Social Sciences.” Working Paper, ArXiv:2203.02198v1.

Rasmussen, SHR., Weinschenk, A., Norgaard, AS., von Bornemann Hjelmborg, J., Klemmensen, R. (2021). “Educational Attainment Has a Causal Effect on Economic, But Not Social Ideology: Evidence from Discordant Twins.” Political Studies, https://doi.org/10.1177/00323217211008788.

Romer, T. (1975). “Individual welfare, majority voting, and the properties of a linear income tax.” Journal of Public Economics, 4(2).

Saribay, SA. and Yilmaz, O. (2017). “Analytic cognitive style and cognitive ability differentially predict religiosity and social conservatism.” Personality and Individual Differences, 114.

Schoon, I., Cheng, H., Gale, CR., Batty, GD. and Deary, IJ. (2010). “Social status, cognitive ability, and educational attainment as predictors of liberal social attitudes and political trust.” Intelligence 38.

Shanahan, MJ., Hofer, SM. (2005). Social Context in Gene–Environment Interactions: Retrospect and Prospect. The Journals of Gerontology: Series B, Special issue 1.

Sidanius, J. (1985). Cognitive Functioning and Sociopolitical Ideology Revisited. Political Psychology, 6(4).

Spinath, FM. and Bleidorn, W. (2017). “The New Look of Behavioral Genetics in Social Inequality: Gene-Environment Interplay and Life Chances.” Journal of Personality, 85(1).

Sterling, K., Jost, JT., Pennycook, K. (2016). “Are neoliberals more susceptible to bullshit?.” Judgment and Decision Making, 11(4).

Trejo, S. and Domingue BJ. (2018). “Genetic nature or genetic nurture? Introducing social genetic parameters to quantify bias in polygenic score analyses.” Biodemography and social biology, 64(3-4).

Turkheimer, E., Harden, KP., D’Onofrio, B., Gottesman, II. (2011). “The Scarr-Rowe Interaction between Measured Socioeconomic Status and the Heritability of Cognitive Ability.” In McCartney, K. and Weinberg, RA. (eds.) Experience and Development, Psychology Press.

van der Waal, J., Achterberg, P. and Houtman, D. (2007). “Class Is Not Dead—It Has Been Buried Alive: Class Voting and Cultural Voting in Postwar Western Societies (1956–1990).” Politics & Society, 35(3).

Weakliem, DL. (2002). “The Effects of Education on Political Opinions.” International Journal of Public Opinion Research 14(2):141–157.

Zagai, U., Lichtenstein, P., Pedersen, NL., Magnusson, PKE. (2019). “The Swedish Twin Registry: Content and Management as a Research Infrastructure.” Twin Research and Human Genetics, 22(6).

Zhang, X. and Belsky, J. (2020). “Three phases of Gene x Environment interaction research: Theoretical assumptions underlying gene selection.” Development and Psychopathology, pp.1–12.

